# Improving the time and space complexity of the WFA algorithm and generalizing its scoring

**DOI:** 10.1101/2022.01.12.476087

**Authors:** Jordan M. Eizenga, Benedict Paten

## Abstract

**Motivation:** Modern genomic sequencing data is trending toward longer sequences with higher accuracy. Many analyses using these data will center on alignments, but classical exact alignment algorithms are infeasible for long sequences. The recently proposed WFA algorithm demonstrated how to perform exact alignment for long, similar sequences in *O*(*sN*) time and *O*(*s*^2^) memory, where *s* is a score that is low for similar sequences (Marco-Sola *et al.*, 2021). However, this algorithm still has infeasible memory requirements for longer sequences. Also, it uses an alternate scoring system that is unfamiliar to many bioinformaticians.

**Results:** We describe variants of WFA that improve its asymptotic memory use from *O*(*s*^2^) to *O*(*s*^3/2^) and its asymptotic run time from *O*(*sN*) to *O*(*s*^2^ + *N*). We expect the reduction in memory use to be particularly impactful, as it makes it practical to perform highly multithreaded megabase-scale exact alignments in common compute environments. In addition, we show how to fold WFA’s alternate scoring into the broader literature on alignment scores.

**Availability:** All code is publicly available for use and modification at https://github.com/jeizenga/wfalm.

**Supplementary information:** Supplementary data are available online.

## 1 Introduction

Pairwise sequence alignments are at the heart of many genomic analyses, from read mapping to genome assembly to genome comparison. Accordingly, these analyses fundamentally require efficient alignment algorithms in order to be practical. This is particularly true for longer sequences, such as assembled contigs or the long reads generated by the modern sequencing platforms produced by PacBio and Oxford Nanopore Technologies. Classical algorithms like Needleman-Wunsch (Needleman and Wunsch, 1970) and Smith-Waterman-Gotoh (Smith and Waterman, 1981; Gotoh, 1982) require quadratic time and memory in the length of the sequences. For longer sequences, this quickly becomes infeasible.

The recently proposed WFA algorithm (Marco-Sola *et al.*, 2021) represents a major step forward in pairwise alignment. Unlike previously proposed acceleration techniques for long sequences, WFA guarantees optimal alignments. However, its run time is bounded by *O*(*sN*), where *N* is the length of the shorter sequence and s is a score that is smaller for more similar sequences. This means that the alignment is feasible even for very long sequences so long as they are highly similar, which is often the case in the applications that interest genomics researchers. In addition, sequences that are relatively high entropy will often require far less time than than the asymptotic upper bound suggests.

Algorithmically, WFA is heavily based on the *O*(*ND*) difference algorithm by Myers (1986). WFA consists mainly of an adaptation of Gotoh’s affine gap technique (Gotoh, 1982) to Myers’ algorithm. In addition, the inventors identified that WFA’s iteration structure allowed modern compilers to automatically apply the SIMD vectorization techniques that many earlier approaches leaned on for efficiency (Farrar, 2007; Suzuki and Kasahara, 2018). These techniques are otherwise highly laborious and error prone to implement by hand.

Certain limitations still frustrate WFA’s applicability in some settings. For one, with sufficiently long or divergent sequences, WFA’s *O*(*s*^2^) memory requirements become a bottleneck before its time requirements. Moreover, such sequences are now routinely produced by increasingly long sequencing reads (>1 Mbp) (Jain *et al.*, 2018) and the increasingly contiguous genome assemblies (>10 Mbp) that these long reads have enabled (Cheng *et al.*, 2021; Shafin *et al.*, 2020).

Another annoyance is that WFA uses an alignment scoring system that is different than conventional algorithms. This makes it inaccessible by the rich literature on probabilistic interpretations of alignments and alignment scores, which is rooted in the conventional scoring scheme (Henikoff and Henikoff, 1992; States *et al.*, 1991; Durbin *et al.*, 1998). Accordingly, it lacks the capacity to use this literature to guide parameterization and interpretation.

In this work, we present results that attenuate these limitations. We propose two alterations to the WFA algorithm that reduce the asymptotic complexity of both its space and its time requirements, respectively. These alterations can be used together or separately. In addition, we prove equivalence results for the WFA scoring system that connect it to classical alignment algorithms. Finally, we demonstrate the improved practicality of the new WFA variants on real genomic sequence data.

## 2 Algorithm

### 2.1 The WFA algorithm

WFA computes a global alignment of two sequences *t* and *q* of length *M* and *N*, respectively. For any particular alignment, define the following variables:

- *X* the total length of mismatches
- *O* the number of insertions and deletions
- *E* the total length of insertions and deletions

Then alignment computed by WFA minimizes the score function

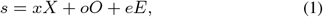

where *x, o*, and *e* are the mismatch, gap open, and gap extend penalties, respectively.

To accomplish this, WFA uses dynamic programming over increasing values of *s* until completing an alignment. In each iteration of *s*, it iterates over a range of values *d* = -*s*,...,*s* that correspond to diagonals in the dynamic programming matrix of conventional alignment algorithms The set of dynamic programming entries for one value of *s* are called a *wavefront* (Figure 1).

**Fig. 1.**
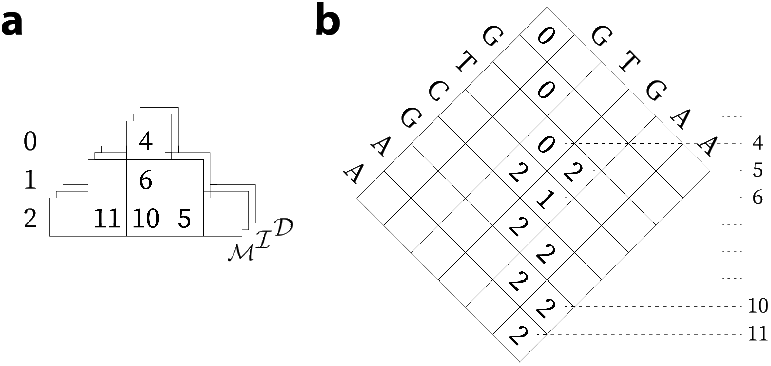
Diagram of the WFA algorithm. The dynamic programming data structure of WFA. (a) is juxtaposed with the dynamic programming matrix used in conventional alignment (b). In this instance, the WFA parameters *x, o,* and e are all 1. Columns in (a) correspond to diagonals in (b), and the matrix is rotated to highlight the correspondence. Rows in (a) are wavefronts for a given score of *s*. The values stored in dynamic programming in (a) correspond to antidiagonals in (b). WFA maintains three parallel dynamic programming structures: 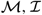, and 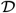. The 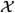 values from Equation 2 are only needed temporarily and are thus omitted.

The dynamic programming recursions are as follows (but note that this formulation differs in minor details from the original published formulation, chiefly that we track antidiagonal indexes rather than row indexes). Let *μ*(*i, j*) be the length of the longest matching prefix between *t_i:M_* and *q_j:N_*, then

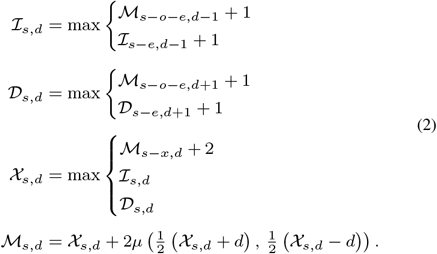

This recurrence begins with the base case 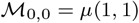, and it continues until 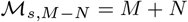.

#### 2.1.1 Complexity analysis

The time complexity of WFA is *O*(*sN*), taking *N* = min{*M, N*} without loss of generality. This time is dominated by computing the *μ* function. Within each of the *O*(*s*) diagonals, it may need to perform *O* (*N*) character comparisons over the course of the algorithm. Also, the algorithm requires *O*(*s*^2^) additional space over and above the *O*(*M* + *N*) space of the input data. This is driven by maintaining the dynamic programming structure in memory so that the alignment can be identified by traceback.

### 2.2 A new variant of WFA with *O*(*s*^3/2^) memory use

With sufficiently long or sufficiently divergent sequences, it is entirely possible that WFA’s *O*(*sN*) run time is practical in real world computing environments while its *O*(*s*^2^) memory footprint is not. In this section, we present a variation on WFA that reduces its additional memory usage to *O*(*s*^3/2^).

The strategy—inspired by WhatsHap (Patterson *et al.*, 2015)—is to retain only a subset of wavefronts in memory after computing them. This subset will contain insufficient information to perform traceback after completing dynamic programming. Thus, blocks of missing wavefronts between the retained wavefronts will be recomputed as needed, used for traceback, and then discarded. This results in each wavefront being computed at most two times, and therefore the run time increases only by a constant factor. The maximum memory usage can be split into two components: 1) the memory of the retained wavefronts, and 2) the largest block of recomputed wavefronts.

#### 2.2.1 A schedule of subsampled wavefronts

In order to be able to recompute non-retained wavefronts, it is necessary to have access to the previous *p* = max{*x, o* + *e*} wavefronts. Accordingly, the wavefronts will be subsampled in units of p consecutive wavefronts, which we refer to as a *stripe*. We propose a schedule that is divided into *epochs* of increasing size, with the *k*-th epoch consisting of 4^*k*^ stripes. Within the *k*-th epoch, we retain every 2^*k*^-th stripe (Figure 2). This strategy allows the schedule of retained wavefronts to be determined online without knowing s in advance.

**Fig. 2.**
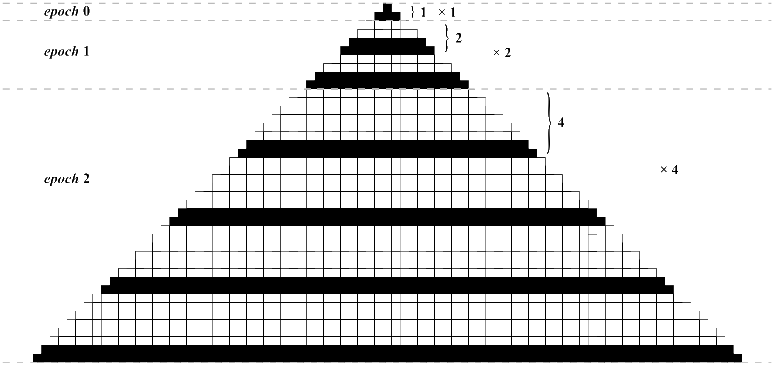
Example of stripe sampling schedule. In this instance, the stripe width *p* is 2. The schedule is divided into epochs. The *k*-th epoch consists of 4^*k*^ stripes, of which we sample every 2^*k*^-th stripe.

##### Theorem 1.

*For constant p, the proposed schedule of retained wavefronts leads to O*(*s*^3/2^) *memory use*.

Proof. We will separately consider the two sources of memory use: the retained wavefronts and the largest block of recomputed wavefronts.

##### Retained wavefronts

At the end of the *k*-th epoch, the total number of stripes is *O*(4^*k*^). Thus, if the WFA dynamic programming terminates in the *n*-th epoch, then *n* = log_4_ *s* + *O*(1). We can use the memory that would be consumed by the entirety of *n* full epochs as an upper bound on the realized memory usage. Since the largest of the 2^*k*^ stripes in the *k*-th epoch covers *O*(4^*k*^) diagonals, we obtain the upper bound

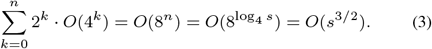

##### Largest block of recomputed wavefronts

The final block of recomputed wavefronts is the largest one. As previously, it occurs in the *n*-th epoch where *n* = log_4_ *s* + *O*(1). The block consists of 2^*n*^ stripes, the widest of which covers *O*(4^*n*^) diagonals. Thus, the total memory use is *O*(8^*n*^) = *O*(*s*^3/2^).

#### 2.2.2 Optimality of the wavefront sampling schedule

One might wonder if it possible to use the same strategy but choose a better schedule of wavefronts to sample. As it turns out, the schedule we have proposed is optimal up to factors of *o*(*s^ε^*) for any *ε* > 0.

##### Theorem 2.

*In the proposed algorithm, there is no schedule of sampled wavefronts that achieves O*(*s*^3/2-*ε*^) *memory usage for any ε* > 0.

Proof. Suppose that the memory usage of the largest block of non-retained stripes requires *O*(*s^α^*) memory to recompute. Further, let the ordinal values of the stripe that ends the *k*-th block be denoted *b_k_*. The memory usage of the *k*-th block is then 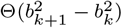. This gives the recursion 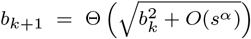 with base case *b*_0_ = 0. By induction, this implies 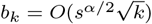.

We turn now to the memory usage of the retained stripes, and our goal will be to provide a lower bound. Accordingly, we consider the selection of retained stripes that will minimize their memory usage. Since the memory usage of the *k*-th stripe is strictly increasing in b_k_, the memory usage is minimized by making each *b_k_* as small as possible. However, this is subject to the constraint that 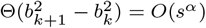. This yields the recursion 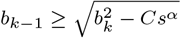 for some constant *C* with base case *b_n_* = *s*, where *n* is the total number of retained stripes. By induction, this implies 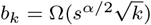 and hence 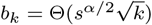.

Using the previous results, we obtain 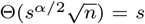, which gives 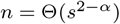. Noting that the memory usage of the *k*-th stripe is Θ(*b_k_*), the total memory of the retained stripes is then given by

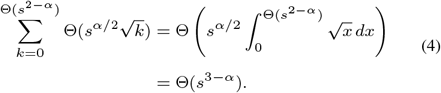

Thus, if 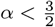 so that the memory required to recompute a non-retained block is *O*(*s*^3/2-*ε*^), then the memory use of the retained stripes is Ω(*s*^3/2+*ε*^).

### 2.3 Achieving *O*(*s*^2^ + *M* + *N*) run time in WFA

WFA’s run time is dominated by the time it takes to compute the *μ* function from Equation 2. In the original implementation, it potentially needs to make *O*(*N*) character comparisons over *O*(*s*) diagonals. In this section, we describe an alternative method to compute the *μ* function in *O*(1) time using *O*(*M + N*) preprocessing. This brings the total run time to *O*(*s*^2^ + *M + N*).

The basic insight for this algorithmic technique was already present in Myers’ *O*(*ND*) publication (1986). However, it was reported as requiring *O*((*M + N*) log(*M + N*)) preprocessing time, which was state-of-the-art when the paper was published. Since then, there have been breakthroughs in data structures that allow linear time.

#### Theorem 3.

*The μ function can be computed in O*(1) *time given O*(*M + N*) *preprocessing time and O*(*M + N*) *additional space*.

Proof. We begin by building a suffix tree of the string *q*$_1_*t*$_2_, which can be done in *O*(*M + N*) time and space (Ukkonen, 1995). In this tree, *μ*(*i,j*) corresponds to the depth (measured in sequence characters) of the lowest common ancestor of the leaves corresponding to *t*_*i:M*_ and *q_j:N_*. The internal nodes of the suffix tree can easily be annotated with their depth after construction. Moreover, there exist techniques to compute lowest common ancestor in a static rooted tree in *O*(1) time using *O*(*M + N*) additional space and preprocessing time (Schieber and Vishkin, 1988).

This brings the total run time per cell in the dynamic programming structure down to *O*(1). Thus, the overall run time of the algorithm becomes *O*(*s*^2^ + *M + N*). The additional space required by the algorithm increases from *O*(*s*^2^) to *O*(*s*^2^ + *M + N*). However, the overall space was already *O*(*s*^2^ + *M + N*), because the sequences had to be kept in memory to perform character comparisons.

### 2.4 Relationship between WFA scoring and conventional alignment scores

The alignments produced by WFA can intuitively be recognized as meaningful alignments. However, the relationship between their scoring and the scoring of conventional alignment algorithms is less immediately apparent. This is unfortunate, because there has been a large amount of research into probabilistic interpretations of the conventional scores. This research can aid in choosing appropriate parameters (Henikoff and Henikoff, 1992; States *et al.*, 1991) or in interpreting scores (Durbin *et al.*, 1998). For local alignments, there are also tractable tests of statistical significance (Karlin and Altschul, 1990).

In this section, we will use a subscript of *w* to denote scores and parameters of <u>WF</u>A and a subscript of *c* to denote scores and parameters of conventional algorithms. We will also add a variable to the list of variables used to describe an alignment in Section 2.1:

- *L* the total length of matches

The score of a conventional alignment can then be expressed

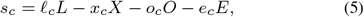

where *ℓ_c_* is the match bonus, and *x_c_, o_c_*, and *e_c_* are the mismatch, gap open, and gap extend penalties, respectively. The objective of conventional algorithms is to find the global or local alignment that maximizes *s_c_*.

#### 2.4.1 Equivalence of conventional global alignment and WFA

The differences between the objective functions in conventional global alignment and WFA are ultimately superficial. For a given set of conventional global alignment parameters, let the WFA parameters be

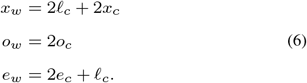

With these parameters and the identity 2*L* + 2*X* + *E* = *M* + *N*, simple algebraic manipulations give

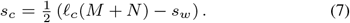

Accordingly, any global alignment that minimizes *s_w_* also maximizes *s_c_* and vice versa. Further, it is possible to choose conventional alignment parameters from a given set of WFA parameters with the converse property.

#### 2.4.2 Optimal seeded local alignment using WFA

While WFA can readily replace the conventional global alignment algorithm through reparameterization, the same is not true for conventional local alignment. We identify and address three obstacles standing in the way.

First, WFA cannot explore all possible pairs of initial characters for a local alignment without losing its run time of *O*(*sN*) or *O*(*s*^2^ + *M + N*) (assuming *N* = min {*M, N*}, see Section 2.3). We address this obstacle by considering a restricted problem: the optimal seeded local alignment. In this variant, we seek the optimal alignment between prefixes of *q* and *t* rather than arbitrary substrings. This problem is of practical importance for alignment in the seed-and-extend methodology, which is widely used in efficient alignment tools (Altschul *et al.*, 1997; Li and Durbin, 2009). We also note that a further variation using suffixes instead of prefixes is trivially similar, so we omit it to streamline the presentation.

The second obstacle is that reducing the number of parameters from four to three (Equation 6) obscures the relative balance between the match bonus and the penalty parameters. This makes it more challenging to differentiate between positive and negative scores, which in turn makes it more challenging to decide when an alignment is optimal during dynamic programming. However, it is still possible to recover the score of aligned prefixes using Equation 7. Note that this equation still holds if we take *M* and *N* to be the length of two prefixes. Moreover, *M + N* is precisely the antidiagonal index, which is what is stored in the WFA dynamic programming structure 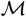. Thus, the score of the corresponding local alignment is 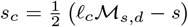, and we begin traceback at the *s* and *d* that maximize this score.

The third and final obstacle is that WFA’s time and memory requirements are bounded by the global alignment score *s_w_*. This score can be large even when there is a high quality local alignment. We provide a technique to partially address this problem. Without loss of generality, we assume *N* = min{*M, N*}, in which case a trivial upper bound on the seeded local alignment score is *ℓ_c_N*. If during the course of WFA, we have computed a dynamic programming value 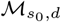, then the trivial bound implies that the optimal seeded local alignment corresponds to a WFA alignment score *s_w_* where 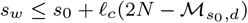. This can be used to limit the total number of iterations in WFA’s outer loop over *s*. This bound is of particular interest when 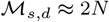, i.e. when there is a nearly full-length local alignment. Accordingly, we expect this condition to be most useful in read mapping, where it is often met.

## 3 Implementation

We have implemented all of the algorithms described in this publication in C++. The code is publicly available for use and modification at https://github.com/jeizenga/wfalm. The implementation depends on SDSL (Gog *et al.*, 2014) for its implementation of a suffix tree. This dependency can be easily omitted if the *O*(*s*^2^ + *M + N*) algorithms are not required. This repository also includes the script used for producing the benchmarking results that follow.

## 4 Methods

We evaluated the run time and memory usage of the standard WFA algorithm and our proposed alterations on two data sets, both generated by the Human Pangenome Reference Consortium (Miga and Wang, 2021). The first set consists of long reads sequenced on the Oxford Nanopore Technologies (ONT) PromethION platform. The second consists of assembled contigs. In both cases, the data are derived from the human individual HG002, and they are subsetted to chromosome 12. For the long reads, the target sequences for the alignment were retrieved by mapping the reads to the assembled contigs with Winnowmap (Jain *et al.*, 2020) and then retrieving the contig sequences. For the contigs, the target sequences were retrieved by mapping the contigs to the T2T Consortium’s v1.1 genome assembly of the haploid CHM13 cell line (Nurk *et al.*, 2021), once again using Winnowmap. Alignment parameters were derived using the method of States *et al.* (1991), assuming sequence identity of 95% for ONT data and 99.9% for the assembled contigs.

We performed all evaluations on AWS r6i.2xlarge instances, which have 64 GB of RAM. Peak memory use was measured with the Unix time utility. Time was measured within the benchmarking script using timing functions from the C++ standard library.

## 5 Results

In both the ONT and contig data, we found that the suffix tree-based algorithm required significantly more time than the direct comparison algorithm. This contrasts with the suffix tree algorithm’s favorable asymptotic time complexity, suggesting that these sequences are insufficiently long or insufficiently divergent for the asymptotic behavior to set in. We exclude the suffix tree algorithm from results figures to make visual comparison easier. Figures that include the suffix tree algorithm are included as Supplementary Figures 1 and 2.

The low memory variant required starkly less memory than standard WFA (Figure 3). In both data sets, a few sequences caused the standard WFA algorithm to exhaust the 64 GB of available RAM. The low memory WFA algorithm used a maximum of 3.7 GB of RAM on any sequence. In 88% of the sequences, it used less than 0.5 GB of RAM.

**Fig. 3.**
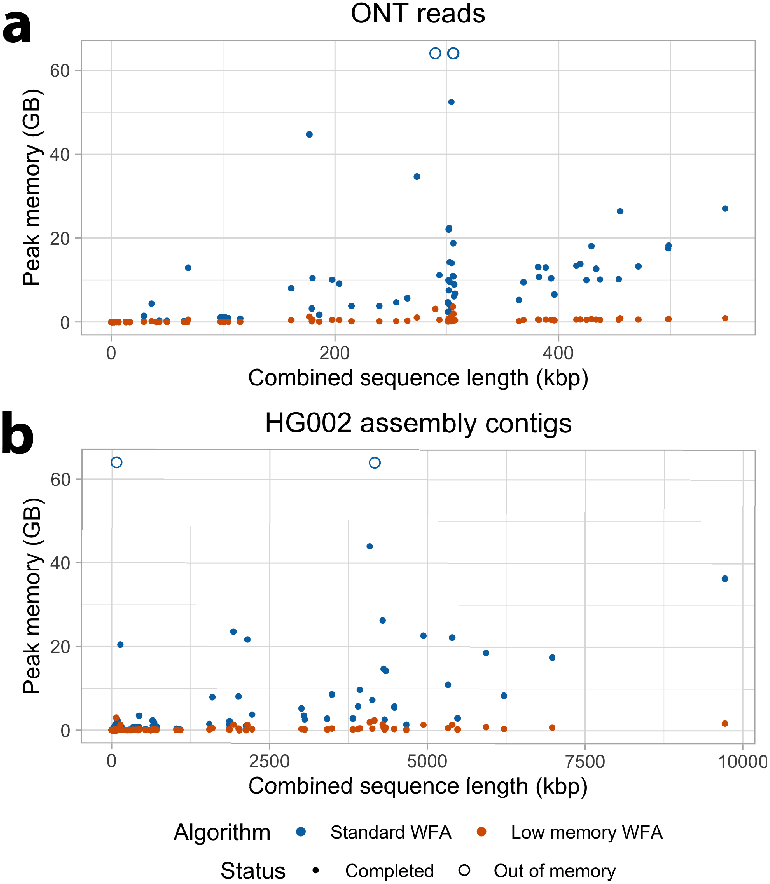
Memory used by the standard *O*(*s*^2^) WFA algorithm and the low memory *O*(*s*^3/2^) WFA variant. Memory is plotted against the sum of the length of the sequences being aligned. Panels show results for ONT reads (a) and assembled contigs (b). If an alignment problem did not complete because it exhausted the 64 GB RAM available, it is shown as an empty circle.

Consistent with expectations, our low memory variant of WFA requires a similar but somewhat greater amount of time to standard WFA, presumably driven by the need to recompute some wavefronts (Figure 4). The ratio between their running times was 1.76 onaverage. Both algorithms were able to align most ONT reads > 100 kbp and assembled contigs > 1 Mbp in a few minutes.

**Fig. 4.**
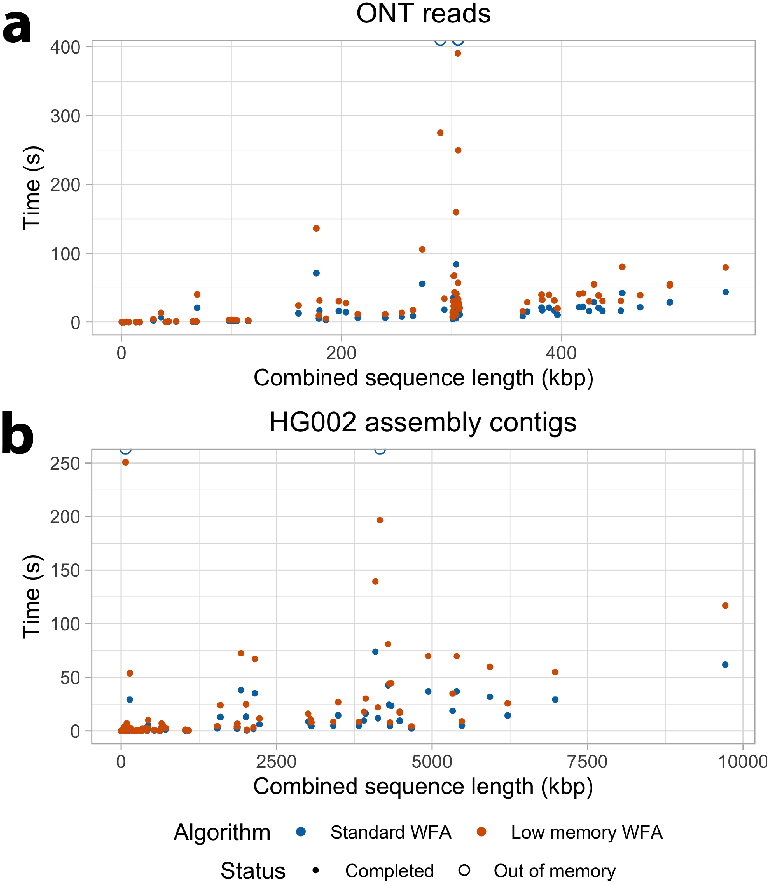
Time used by the standard *O*(*s*^2^) WFA algorithm and the low memory *O*(*s*^3/2^) WFA variant. Time is plotted against the sum of the length of the sequences being aligned. Panels show results for ONT reads (a) and assembled contigs (b). If an alignment problem did not complete because it exhausted the 64 GB RAM available, it is shown as an empty circle.

## 6 Discussion

Future sequencing data present a challenge to current algorithmic methods. The technology is trending toward producing longer reads at higher quality. This in turn is enabling highly contiguous genome assemblies, which are driving developments in pangenomics, metagenomics, and comparative genomics. Accordingly, we expect that both long read resequencing and whole genome comparison will be increasingly important methodologies moving forward. The scale and characteristics of these data tax current methods, and algorithms must adapt.

In this paper, we have described extensions to the recently introduced WFA algorithm, which can efficiently compute optimal alignments for highly similar sequences. Two of our extensions reduce its asymptotic complexity. The first reduces the space from *O*(*s*^2^) to *O*(*s*^3/2^). The second reduces the time from *O*(*sN*) to *O*(*s*^2^ + *M + N*). In addition, we describe techniques to fold WFA into the broader literature on alignment scoring.

For the time being, the improvements to asymptotic run time have not translated into improvements in actual run time. This concords with conventional wisdom about algorithms based on suffix trees. They frequently have very attractive asymptotic complexity, but the asymptotic analysis conceals large constants in memory and time. In this instance, the memory use was not prohibitive, but the time was. Yet, the suffix tree algorithm may become advantageous in the future with longer sequence lengths. It also may show advantages in sequences that have high levels of internal repetition, such as satellite arrays. In these sequences, the higher rate of spuriously matching sequence makes the direct comparison approach comparatively more costly.

In the case of the low memory WFA variant, we expect that our proposed algorithm will be immediately useful. For the first time, it makes it feasible to compute optimal alignments of megabase long sequencing reads (or similarly diverged sequences) in a couple of GB of RAM. This will make it possible to perform many of these alignments simultaneously in a multithreaded compute server with typical specifications. Nearly all modern tools for efficient sequence analysis require such multithreading to be practical. With the benchmarking data, we can coarsely estimate that, using 16 threads, the low memory WFA can align about 50 kbp/s of ONT data or 727 kbp/s of assembly contig data using a maximum of about 20 GB of RAM (although our data selection procedures probably bias this estimate somewhat). Contrast this to the standard WFA algorithm, which could not align the benchmarking data with 64 GB of RAM using a single thread.

Alignment is a core component of many analyses in genomics, but it is rarely sufficient to perform a complete analysis in itself. The algorithms we have described will need to be integrated into more fully-featured tools to have their full impact. Further, WFA natively produces global alignments, but in many use cases global alignment is not what is required. Resequencing experiments typically perform local or semiglobal alignments. Whole genome comparisons must additionally contend with noncollinear sequence relationships arising from translocation, inversion, gene conversion, and segmental duplication. In this work, we do describe a method to use WFA as an engine for computing local alignments, but these results come with more caveats than other areas of this paper. For these reasons, we fully expect that it will continue to be necessary to use some approximate methods in these tools. Nevertheless, our improvements to WFA will make it possible lean on approximate alignment to a much lesser extent while remaining efficient at the frontiers of genome-scale data.

## Acknowledgements

We would like to thank the Human Pangenome Reference Consortium for sharing a subset of the data they have generated with us for the purpose of benchmarking.

## References

Altschul, S. F. et al. (1997). Gapped BLAST and PSI-BLAST: a new generation of protein database search programs. Nucleic acids research, 25(17), 3389–3402.

Cheng, H. et al. (2021). Haplotype-resolved de novo assembly using phased assembly graphs with hifiasm. Nature Methods, 18(2), 170–175.

Durbin, R. et al. (1998). Biological sequence analysis: probabilistic models of proteins and nucleic acids. Cambridge University Press.

Farrar, M. (2007). Striped Smith–Waterman speeds database searches six times over other simd implementations. Bioinformatics, 23(2), 156–161.

Gog, S. et al. (2014). From theory to practice: Plug and play with succinct data structures. In International Symposium on Experimental Algorithms, pages 326–337. Springer.

Gotoh, O. (1982). An improved algorithm for matching biological sequences. Journal of Molecular Biology, 162(3), 705–708.

Henikoff, S. and Henikoff, J. G. (1992). Amino acid substitution matrices from protein blocks. Proceedings of the National Academy of Sciences, 89(22), 10915–10919.

Jain, C. et al. (2020). Weighted minimizer sampling improves long read mapping. Bioinformatics, 36(Supplement_1), i111–i118.

Jain, M. et al. (2018). Nanopore sequencing and assembly of a human genome with ultra-long reads. Nature Biotechnology, 36(4), 338–345.

Karlin, S. and Altschul, S. F. (1990). Methods for assessing the statistical significance of molecular sequence features by using general scoring schemes. Proceedings of the National Academy of Sciences, 87(6), 2264–2268.

Li, H. and Durbin, R. (2009). Fast and accurate short read alignment with Burrows–Wheeler transform. Bioinformatics, 25(14), 1754–1760.

Marco-Sola, S. et al. (2021). Fast gap-affine pairwise alignment using the wavefront algorithm. Bioinformatics, 37(4), 456–463.

Miga, K. H. and Wang, T. (2021). The need for a human pangenome reference sequence. Annual Review of Genomics and Human Genetics, 22.

Myers, E. W. (1986). An O(ND) difference algorithm and its variations. Algorithmica, 1(1-4), 251–266.

Needleman, S. B. and Wunsch, C. D. (1970). A general method applicable to the search for similarities in the amino acid sequence of two proteins. Journal of Molecular Biology, 48(3), 443–453.

Nurk, S. et al. (2021). The complete sequence of a human genome. bioRxiv.

Patterson, M. et al. (2015). WhatsHap: weighted haplotype assembly for future-generation sequencing reads. Journal of Computational Biology, 22(6), 498–509.

Schieber, B. and Vishkin, U. (1988). On finding lowest common ancestors: Simplification and parallelization. SIAM Journal on Computing, 17(6), 1253–1262.

Shafin, K. et al. (2020). Nanopore sequencing and the Shasta toolkit enable efficient de novo assembly of eleven human genomes. Nature Biotechnology, 38(9), 1044–1053.

Smith, T. F. and Waterman, M. S. (1981). Comparison of biosequences. Advances in Applied Mathematics, 2(4), 482–489.

States, D. J. et al. (1991). Improved sensitivity of nucleic acid database searches using application-specific scoring matrices. Methods, 3(1), 66–70.

Suzuki, H. and Kasahara, M. (2018). Introducing difference recurrence relations for faster semi-global alignment of long sequences. BMC Bioinformatics, 19(1), 33–47.

Ukkonen, E. (1995). On-line construction of suffix trees. Algorithmica, 14(3), 249–260.

