## Supplementary Material for "Improving the time and space complexity of the WFA algorithm and generalizing its scoring"

Jordan M. Eizenga
*University of California Santa Cruz Genomics Institute*

Benedict Paten
*University of California Santa Cruz Genomics Institute*

#### 1 Supplementary methods

All custom scripts described in this section are available in the GitHub repository `https://github.com/jeizenga/wfalm`.

##### 1.1 Retrieving paired sequences

The paired sequences for the ONT data alignments were identified with the following `winnowmap` (Jain *et al.*, 2020) script.

```
meryl count k=15 output merylDB HG002.f1_assembly_v2_genbank.dip.fa.gz
meryl print greater-than distinct=0.9998 merylDB > repetitive_k15.txt
winnowmap -W repetitive_k15.txt -a -x map-ont -Y -L --eqx --cs -I8g -t8 HG002.f1_assembly_[illegible]
reads.fq
```

The sequences were extracted by using `samtools view` (Li *et al.*, 2009) chained with `paftools sam2paf` (Li, 2016) to convert the alignments into PAF format. We then filtered out any mappings with MAPQ $<$ 20 or sequence length $<$ 150 kbp. Next, we converted the PAF into BED format with a custom script and extracted the corresponding subsequences from the mapping target sequences using `samtools faidx`.

The paired sequences for the HG002 assemblies were identified with this `winnowmap` script.

```
meryl count k=19 output merylDB chm13.draft_v1.1_plus38Y.fa.gz.fa
meryl print greater-than distinct=0.9998 merylDB > repetitive_k19.txt
winnowmap -W repetitive_k19.txt --MD -a -x asm5 -t32 chm13.draft_v1.1_plus38Y.fa.gz
HG002.f1_assembly_v2_genbank.dip.fa.gz
```

We converted the mappings into PAF format by chaining `samtools view` with `paftools sam2paf`. We then used a custom script to greedily merge adjacent mapped regions of the assemblies. Specfically, we merged adjacent mapped regions if they were consistently oriented to each other on both the CHM13 and HG002 assemblies and were separated by $<$ 500 bp on both sequences. Mappings with MAPQ $<$ 30 were filtered from this step, and merged regions were terminated if a merge would result in a sequence that was > 10 Mbp long. This merged PAF was then converted into two BED files (one for the CHM13 assembly and one for the HG002 assembly), and the sequences were extracted using `samtools faidx`.

##### 1.2 Benchmarking performance

Benchmarking was performed by a C++ benchmarking script that reads sequences from a FASTA file, aligns them with one of algorithms described in this paper, and then exits. It measures its own time using the C++ standard library function `std::chrono::steady_clock` and its baseline memory usage (before entering the algorithm) with the C++ Linux function `getrusage`. Both of these values are printed to logs. This C++ benchmark was run in a loop over different algorithmic options with a Bash script, which also measured maximum memory usage using the Unix `time` utility. The resulting logs were scraped by a custom Python script and converted into a table format, which was used to create figures using `ggplot2` (Wickham, 2011).

#### 2 Supplementary figures

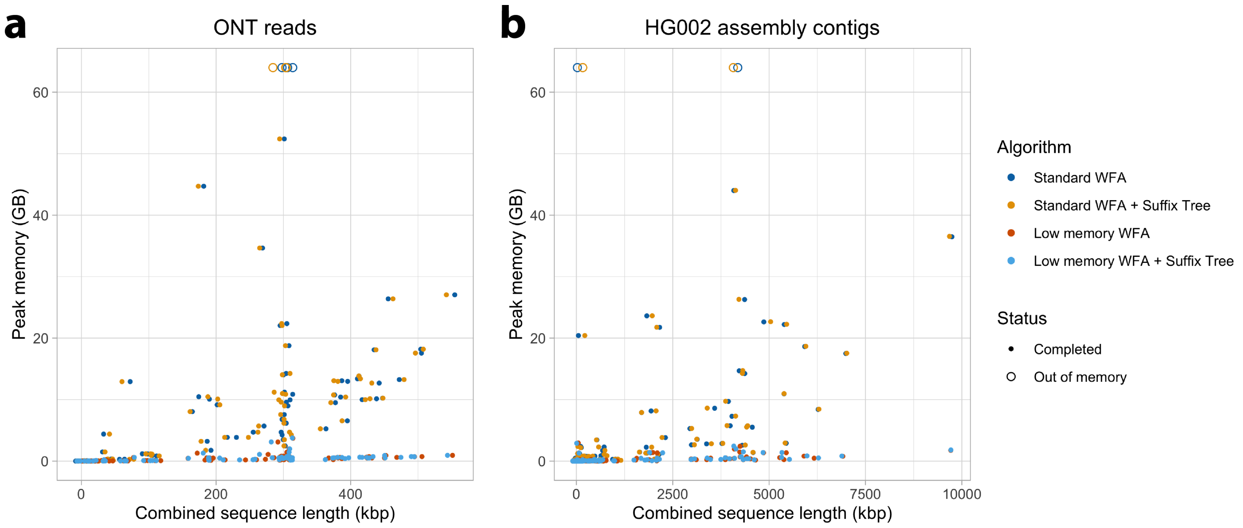


Figure 1: *Memory used by the standard $O(s^2)$ WFA algorithm and the low memory $O(s^{3/2})$ WFA variant, either with or without the $O(s^2 + N + M)$ suffix tree variant.* Memory is plotted against the sum of the length of the sequences being aligned. Panels show results for ONT reads (**a**) and assembled contigs (**b**). If an alignment problem did not complete because it exhausted the 64 GB RAM available, it is shown as an empty circle. Random horizontal jitter has been added to make points easier to differentiate (with jitter values in $[-10, 10]$ for ONT and $[-100, 100]$ for the HG002 assembly).

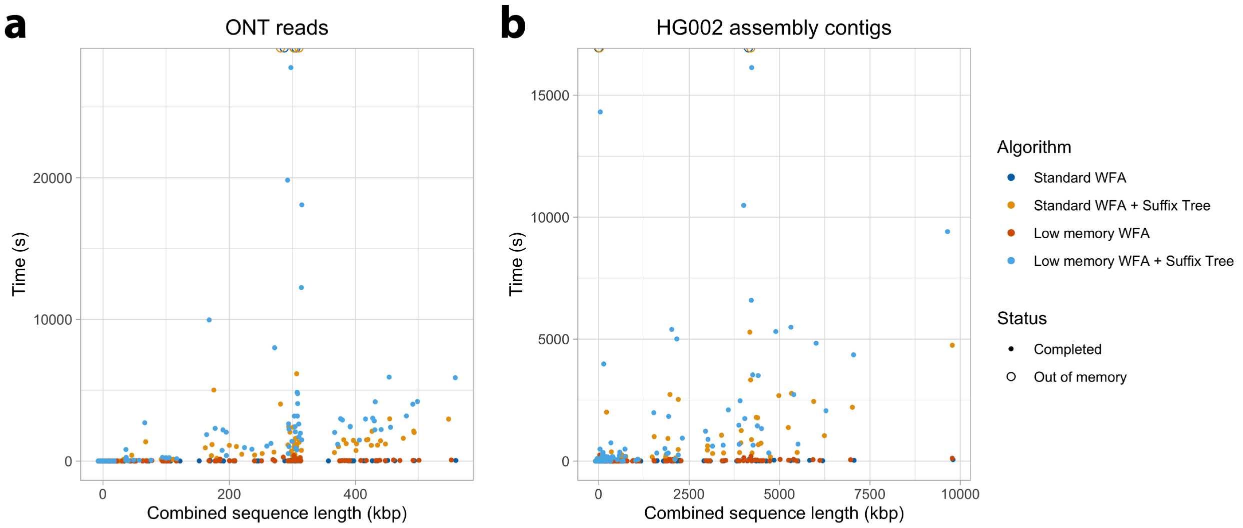


Figure 2: *Time used by the standard $O(s^2)$ WFA algorithm and the low memory $O(s^{3/2})$ WFA variant, either with or without the $O(s^2 + N + M)$ suffix tree variant.* Time is plotted against the sum of the length of the sequences being aligned. Panels show results for ONT reads (**a**) and assembled contigs (**b**). If an alignment problem did not complete because it exhausted the 64 GB RAM available, it is shown as an empty circle. Random horizontal jitter has been added to make points easier to differentiate (with jitter values in $[-10, 10]$ for ONT and $[-100, 100]$ for the HG002 assembly).

### References


Jain, C. *et al.* (2020). Weighted minimizer sampling improves long read mapping. *Bioinformatics*, **36**(Supplement_1), i111–i118.

Li, H. (2016). Minimap and miniasm: fast mapping and de novo assembly for noisy long sequences. *Bioinformatics*, **32**(14), 2103–2110.

Li, H. *et al.* (2009). The sequence alignment/map format and SAMtools. *Bioinformatics*, **25**(16), 2078–2079.

Wickham, H. (2011). ggplot2. *Wiley Interdisciplinary Reviews: Computational Statistics*, **3**(2), 180–185.
